# From Routine Pathology to Precision Oncology: Automated FFPE Tissue Processing for Large-Scale Molecular Studies

**DOI:** 10.64898/2026.08.12.744404

**Authors:** Jéssica Guedes, Andrzej Sliwa-Gonzalez, Leticia Szadai, Philipp Geiger, Nicole Woldmar, Marisol Ayala Reyes, Ramiro Alonso Bastida, Diana Lashidua Fernández Coto, Henriett Oskolás, Matilda Marko-Varga, Lesley Schultz, Roger Appelqvist, Elisabet Wieslander, Johan Malm, György Marko-Varga, Jeovanis Gil

## Abstract

Melanoma incidence continues to rise globally, with formalin-fixed paraffin-embedded (FFPE) tissue archives representing an invaluable resource for large-scale retrospective proteomic studies. However, inconsistent deparaffinization remains a critical pre-analytical bottleneck limiting protein yield, reproducibility, and downstream data quality. In this study, we developed and validated a fully automated FFPE deparaffinization workflow using the Fluent® 780 liquid handling workstation (Tecan ©) and evaluated its performance against a conventional manual protocol in a cohort of 54 patients with primary cutaneous melanoma, predominantly at early AJCC 8th edition stage I–II. The automated workflow achieved superior protein identification (6,146 ± 860 vs. 4,941 ± 1,091 proteins; p < 0.0001) with lower technical variability, while maintaining highly comparable global proteomic profiles as confirmed by principal component analysis and hierarchical clustering. A total of 8,305 proteins (96.1%) were identified by both methods, supporting the reproducibility and equivalence of the automated approach. Patients were stratified by the presence (N=21) or absence (N=33) of histological regression in the primary tumor. Proteomic comparison revealed 97 upregulated and 226 downregulated proteins in regressing melanomas, with pathway enrichment analysis demonstrating elevated mitochondrial and translational activity alongside reduced innate immune and complement pathway activation in the regression group. No statistically significant differences in overall, disease-free, or progression-free survival were observed between groups, consistent with the early-stage composition of the cohort. Digital pathology validated tissue morphology preservation across processing conditions. These findings support the integration of automated FFPE processing with proteomic and digital pathology workflows as a scalable platform for precision melanoma research.

**TOC Figure:** 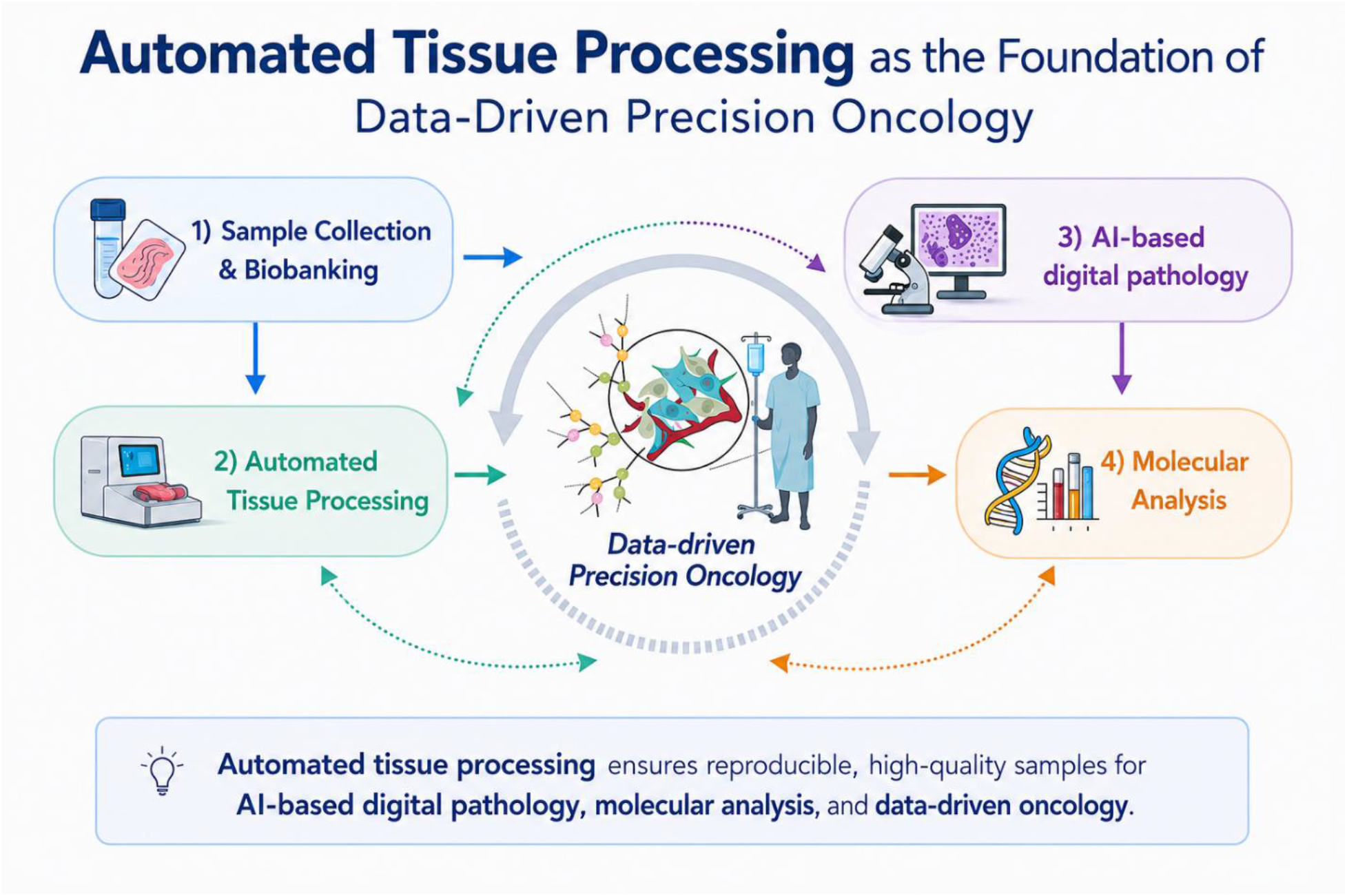

## 1. INTRODUCTION

In recent years, melanoma incidence has shown a steady increase across the European Union, particularly in fair-skinned populations, and accounts for 80% of skin cancer–related deaths worldwide [1–3]. The highest rise is observed among individuals aged 55 and older, likely due to cumulative UV exposure over decades. However, younger adults aged under 50 years, especially women are also seeing rising melanoma rates, possibly linked to indoor tanning and lifestyle factors [4].

Melanoma is a particularly challenging area of oncology, both in terms of disease biology and in the development of effective, patient centric therapies. Among all cancers, melanoma has one of the fastest rates of metastatic spread, often progressing silently through small, undetectable tumor colonies until late stages. These metastases are frequently missed during routine monitoring, making early detection and precision diagnostics a critical unmet need, especially in advanced stage III and IV patients, where therapeutic windows are limited. Regarding the therapeutical options, one of the frequently applied immunotherapy in metastatic melanoma, pembrolizumab (PD1 inhibitor, Keytruda), costs $204,000 per patient annually [5]. Improving early prediction of recurrence is critical to optimize surveillance strategies and guide potential adjuvant interventions. Therefore, accurate stratification of patients according to risk of progression and likelihood of therapy response is a critical clinical need [6].

To tackle this, large-scale biobanks and national tumor registries are increasingly turning to formalin-fixed paraffin-embedded (FFPE) tumor blocks as a core asset for retrospective and prospective studies. These FFPE archives, currently comprising approximately 6 million tumor tissue blocks within a single hospital in our melanoma network, represent one of the largest clinically annotated pathology repositories available for translational cancer research [7]. As the most abundant and widely accessible form of preserved tumor material in pathology departments worldwide, FFPE biobanks constitute an invaluable resource for large-scale molecular investigations, linking proteogenomics, digital pathology, AI-driven image analysis, and comprehensive clinical follow-up to advance precision oncology.

For patient stratification, the use of FFPE-derived samples combined with proteomic analysis offers a complementary strategy by directly capturing protein-level alterations linked to metabolic reprogramming, immune evasion, and tumor–microenvironment interactions, key processes increasingly associated with early melanoma progression [7,8]. However, unlocking the full potential of FFPE blocks for multi-omics investigations including proteomics, genomics, transcriptomics, and metabolomics, requires the careful reversal of formalin-induced crosslinking and removal of paraffin. Traditional solvent-based protocols increase technical variability and protein loss, while formaldehyde-induced cross-links limit protease accessibility and peptide recovery [9–11]. Thus, the development of standardized, automated sample processing pipelines, using robotic platforms for sectioning, deparaffinization, tissue digestion, and molecular extraction, is essential. Such systems not only reduce variability and sample loss, but also make high-throughput, reproducible analysis feasible across diverse clinical biobanks. They are not toxic and environmental friendly. These innovations open up the way for comprehensive molecular characterization of melanoma, enabling the discovery of early biomarkers, therapeutic targets, and resistance mechanisms that can ultimately inform more personalized treatment strategies.

In this study, we compare a conventional in-house deparaffinization workflow with a standardized, automated protocol (TECAN Fluent Control 780) and evaluate their impact on downstream proteomic profiling. Using a cohort of 56 patients with predominantly thin, early-stage melanomas (AJCC 8th edition stage I–II) [12], we aimed to characterize molecular features associated with non-progressive disease. The automated approach enabled reproducible processing of routinely archived clinical FFPE material, supporting scalable proteomic analysis. Furthermore, integration of digital pathology allowed identification of key histomorphological features of non-progressing tumors, highlighting the potential biological relevance of tumor regression in early-stage melanoma. Importantly, the automated deparaffinization protocol facilitates high-throughput and time-efficient workflow in translational dermatologic research.

## 2. RESULTS

The melanoma tissue workflow is based on a thoroughly optimized and standardized protocol developed for large-scale translational clinical studies, ensuring robust, reproducible, and high-throughput processing of FFPE tumor specimens. Following standardized biobanking and automated tissue processing, samples undergo AI-assisted digital pathology with high-resolution whole-slide imaging for comprehensive histopathological evaluation and precise region-of-interest selection, followed by downstream molecular analyses including genomic, proteomic, and biomarker profiling. This integrated precision oncology platform combines automated tissue processing, digital pathology, multi-omics, and clinical metadata, emphasizing standardized and reproducible workflows to generate high-quality molecular data for investigations of tumor heterogeneity, metastatic evolution, biomarker discovery, and precision oncology **(Figure 1)**.

**Figure 1.**
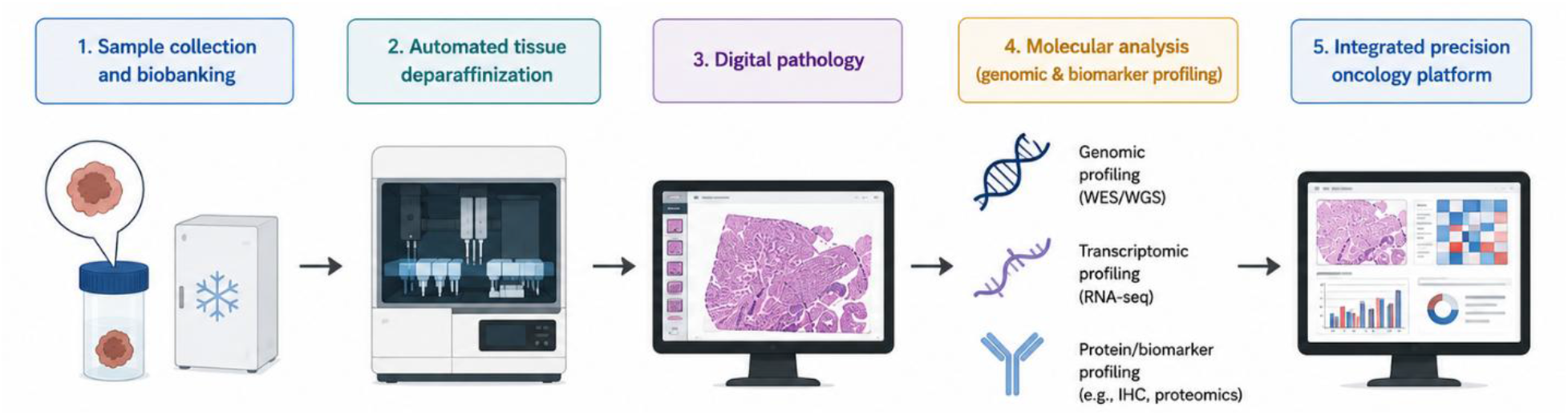
Overview of the automated melanoma tissue processing workflow for integrated digital pathology and molecular analysis.

### 2.1. Clinicopathological characteristics of the study cohort

A clinically well-characterized cohort of 54 patients with primary cutaneous melanoma was established to investigate the biological and molecular significance of histological regression, a key but incompletely understood feature of melanoma progression. Patients were stratified according to the presence (n = 21) or absence (n = 33) of histological regression in the primary tumor, providing the basis for comprehensive clinicopathological, proteogenomic, and biomarker analyses aimed at elucidating the mechanisms underlying melanoma evolution and disease outcome. Patients with regression were older at the time of diagnosis, with a mean age of 69.4 ± 11.6 years, compared to 56.9 ± 19.9 years in the non-regression group. Survival outcomes included disease-free survival (DFS), progression-free survival (PFS), and overall survival (OS). The mean DFS was 90.1 months (±49), the mean PFS was 97.4 months (±40.9), and the mean OS was 97.8 months (±40). Survival data were unavailable for three patients; therefore, these cases were excluded from the survival analyses. No statistically significant differences were observed in disease-free survival (DFS, log-rank *p* = 0.546) and progression-free survival (PFS, log-rank *p*=0.578) between primary melanomas with and without histological regression. However histopathologic regression was associated with improved overall survival in this cohort (log-rank *p* = 0.028), only 18 deaths occurred during follow-up, therefore these results should be considered exploratory and require confirmation in larger studies (**Table 1**).

**Table 1.** Clinicopathological characteristics of patients with primary melanoma with regression and primary melanoma without regression. The table summarizes age at primary melanoma diagnosis, Breslow thickness, sex distribution, AJCC8 clinical stage at diagnosis, histological subtype, and ulceration status in the two study groups. Continuous variables are presented as mean ± standard deviation (SD), while categorical variables are shown as absolute numbers of cases. Abbreviations: AJCC8, American Joint Committee on Cancer, 8th edition; SSM, superficial spreading melanoma; NM, nodular melanoma; LMM, lentigo maligna melanoma.

| Clinicopathological data |  | Primary with regression (N=21) | Primary without regression (N=33) |
| --- | --- | --- | --- |
| Patients (N=54) | Variables | Mean (± SD) | Mean (± SD) |
| Age | Age at primary melanoma diagnosis | 69.4 yrs ($\pm 11.6$) | 56.9 yrs ($\pm 19.9$) |
| Breslow thickness | mean | 0.9 mm ($\pm 0.9$) | 0.92 mm ($\pm 0.89$) |
| Gender | Male | 13 | 13 |
|  | Female | 8 | 12 |
| Clinical stage (AJCC8) at diagnosis | IA | 16 | 19 |
|  | IB - IV | 5 | 14 |
| Histotype of primary melanoma | SSM | 19 | 21 |
|  | **SSM with vertical growth** | 2 | 0 |
|  | **NM** | 0 | 4 |
|  | **LMM** | 0 | 4 |
|  | **Others** | 0 | 4 |
| **Ulceration** | **Yes** | 7 | 12 |
|  | **No** | 14 | 21 |

The mean Breslow thickness was comparable between the groups, measuring 0.9 ± 0.9 mm in melanomas with regression and 0.92 ± 0.89 mm in melanomas without regression. According to the AJCC8 clinical staging system, most patients were diagnosed at stage IA in both groups, 16 in the regression group and 19 in the non-regression group, whereas stages IB–IV were observed in 5 and 14 patients, respectively. Superficial spreading melanoma (SSM) was the predominant histological subtype in both cohorts, accounting for 19 cases in the regression group and 21 cases in the non-regression group. Ulceration was present in 7 patients with regression and in 12 patients without regression, while the absence of ulceration was documented in 14 and 21 cases, respectively. Importantly, the workflow was evaluated across a wide spectrum of primary melanoma tissue architectures, encompassing tumor-rich, tumor-poor, stroma-rich, and immune cell-rich lesions. The successful processing of these morphologically diverse specimens demonstrates the robustness of the automated platform and its capacity to generate high-quality proteomic data irrespective of tumor cellularity, stromal abundance, or immune infiltration (**Figure 2.**)

**Figure 2.**
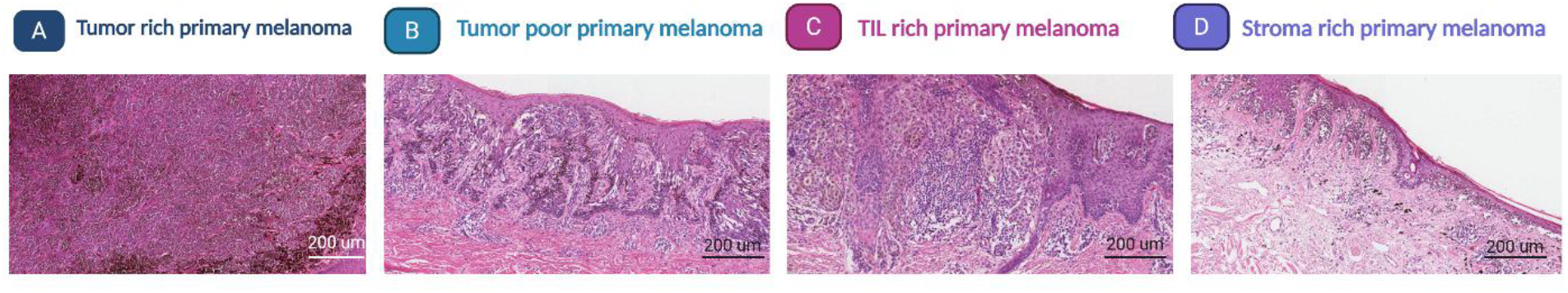
Representative hematoxylin and eosin (H&E)-stained sections illustrating the diversity of tissue architectures encountered across the melanoma cohort. (A) Tumor-rich primary melanoma characterized by high tumor cellularity with minimal stromal content. (B) Tumor-poor primary melanoma exhibiting low tumor cellularity, abundant stromal and TIL rich tissue, and scattered tumor nests. (C) Primary melanoma with prominent tumor-infiltrating lymphocytes (TIL-rich), representing an immune cell-rich tumor microenvironment. (D) Stroma-rich primary melanoma characterized by extensive fibrotic stroma, sparse tumor cells, and low tumor cellularity. Scale bar: 200 μm.

### 2.2. Histomorphological Profiling of FFPE Melanoma Samples by Digital Pathology for automated deparaffinization

From the perspective of a treating clinician, one of the greatest challenges in everyday melanoma care is that while every patient contributes valuable tumor tissue to the pathology archive, only a limited proportion of these specimens can currently be translated into comprehensive molecular information capable of guiding biological understanding and future patient management. Motivated by this unmet clinical need, we developed and validated an automated FFPE tissue processing platform across a broad spectrum of tumor tissue types, establishing a standardized, robust, and scalable workflow that unlocks the molecular potential of routine pathology specimens for large-scale translational studies and precision oncology. This includes samples with varying tumor cell content, stromal composition, and immune cell (leukocyte) infiltration. By encompassing this diversity, the system is designed to accommodate the full range of patient sample variability encountered across cancer studies. This approach enhances robustness, reproducibility, and readiness for clinical implementation in multi-indication oncology settings. To complement the proteomic characterization and validate the suitability of the automated deparaffinization pipeline for downstream histomorphological analysis, digital pathology was applied to H&E-stained whole-slide images of the study cohort. Representative cases were selected to illustrate the full spectrum of tumor tissue heterogeneity encountered across the 54-patient cohort, encompassing samples with high tumor cell content, stroma-dominant architecture, and variable degrees of immune cell infiltration (**Figure 3**).

**Figure 3.**
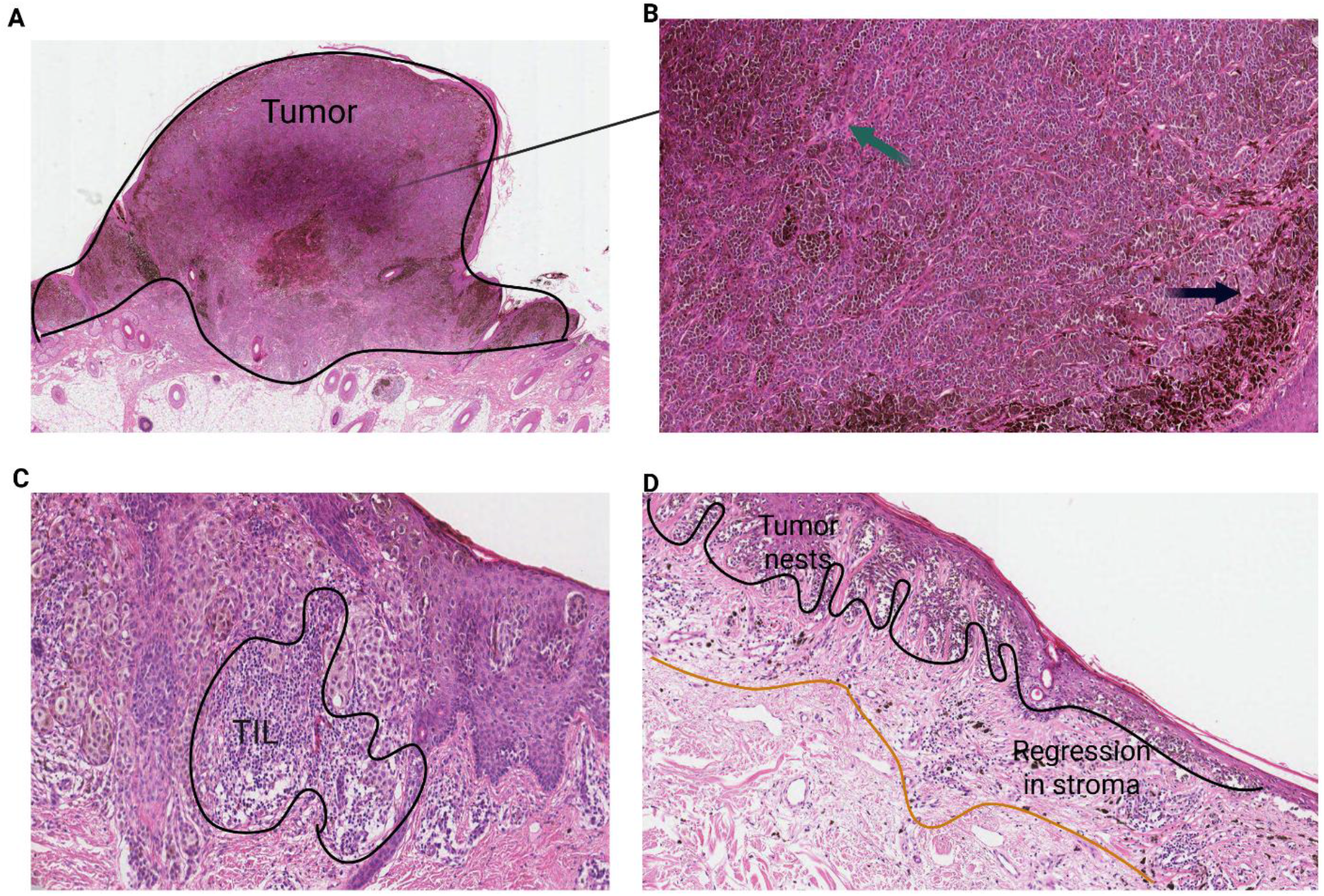
Representative H&E-stained whole-slide image panels illustrating the range of tumor tissue composition across the study cohort. **(A–B)** Samples with high tumor cell content (melanophages as well, marked with black arrow) and minimal stromal (green arrow) involvement (scale bars: **A**, 1000 µm; **B**, 100 µm). **(C)** Case with prominent tumor-infiltrating lymphocyte (TIL) infiltration, including both peritumoral and intratumoral distributions (scale bars: C, 100 µm). **(D)** Case with regression composed of lymphocytes, melanophages and stromal part (orange line) with dispersed tumor nests (black line) (scale bars: 200 µm).

Histopathology annotation of tumor, stromal, and lymphocytic compartments enabled systematic and reproducible characterization of tissue composition across cases. Tumor nests were delineated from surrounding desmoplastic or loose stroma, and tumor-infiltrating lymphocytes (TILs) were assessed both at the tumor periphery and within the tumor parenchyma. Among the cases with histological regression, digital pathology review revealed a characteristic tissue architecture marked by fibrotic stromal replacement, melanophage deposition, and lymphocytic infiltration at the site of regressed tumor (**Figure 2, D**). These features were identified mostly across regression-positive cases regardless of overall tumor cellularity. In contrast, non-regression melanomas displayed more compact tumor nests with variable stromal content. Qualitative assessment of tumor cell content across the cohort demonstrated considerable heterogeneity, ranging from samples with abundant tumor cellularity to those dominated by stromal and inflammatory components. This variability underscores the importance of careful morphological evaluation prior to downstream molecular analyses, and confirms that the automated deparaffinization workflow preserved tissue morphology sufficiently for reliable histopathological interpretation.

### 2.3. Automated Deparaffinization Workflow

Automated deparaffinization was conducted using a Fluent® 780 liquid handling workstation (Tecan, Männedorf, Switzerland) equipped with an 8-channel airFCA pipetting arm (Flexible Channel Arm), an RGA (Robotic Gripper Arm) with centric fingers, four BioShake thermal shakers (Bioshake D30-T EL, QInstruments) with appropriate plate adapters (Adapter-NUNC® & Axygen® deep well 96/2.0 ml, QInstruments), and a temperature-controlled centrifuge (Rotanta 460 RSC, Hettich). The system was programmed to process two microplates in parallel, thereby maximizing throughput and ensuring workflow consistency.

The protocol was adapted from [13], with modifications to enable fully automated paraffin handling. In contrast to the original manual method, where solidified paraffin is removed by hand using a pipette tip, the automated workflow incorporates an additional melting phase, allowing paraffin to be aspirated in its liquid state. While this modification is less efficient at removing paraffin due to residual adherence to well walls, the protocol compensates by increasing the number of extraction cycles. The automated workflow consists of 5–6 repeated cycles of the following steps (see **Figure 4**):

1. **Addition of Envision Buffer:** A variable volume of Envision buffer (400 uL for first round, then 600uL for remaining rounds) is dispensed into each well using a multi-pipetting liquid class.
2. **Heating and Shaking:** Samples are heated at 98 °C and agitated at 900 RPM for 5 minutes to facilitate paraffin melting. 3)**Centrifugation:** Plates are centrifuged at 4°C for 8 minutes at 1000G to separate the melted paraffin from the sample. 4)**Reheating:** Samples are reheated without shaking at 98 °C for 9 minutes to maintain paraffin as the upper liquid phase. 5) **Aspirating Paraffin:** The upper paraffin layer is aspirated using 1000 µL tips (see **Figure 2**). 6) **Tip Disposal:** Tips containing paraffin are discarded. 7) **Cycle Repetition:** Steps 1–6 are repeated for a total of 5–6 cycles to ensure efficient paraffin removal. 8) **Sample Transfer:** After 5–6 cycles, samples were transferred using a 1000 µL wide-bore pipette tip into Azenta 1.0 mL 2D Barcoded Storage Tubes (68-0702, Azenta Life Sciences), held in a 2 mL Axygen® 96-well deep-well plate (P-DW-20-C, Corning Life Sciences) used as a tube rack adapter. One additional deparaffinization cycle was then performed. Finally, samples were transferred to a 1.5 mL microcentrifuge tube (Eppendorf) for further processing. **Liquid Handling Optimization:** To optimize paraffin extraction, custom liquid classes were programmed with the following parameters (see **Figure 5**):

**Figure 4.**
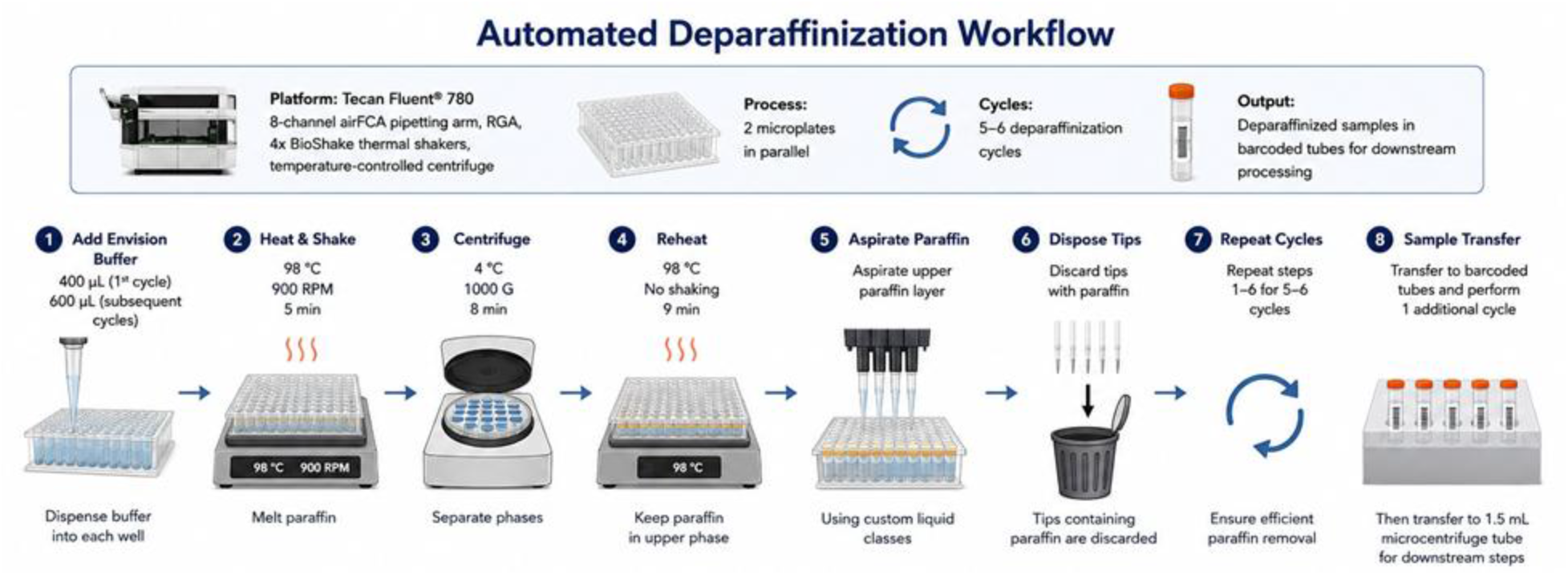
Schematic representation of the automated deparaffinization workflow.

**Figure 5.**
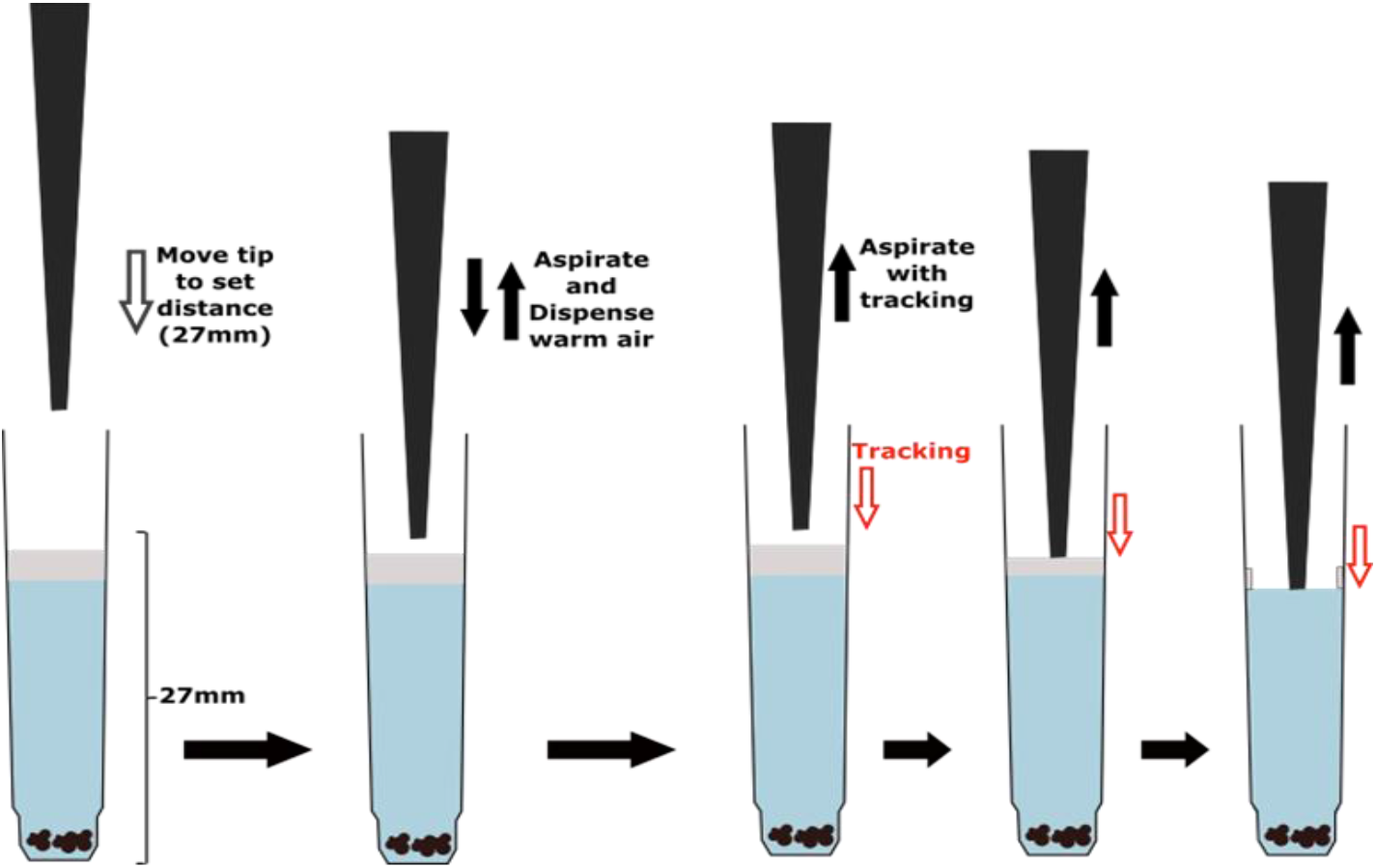
Schematic of the paraffin aspiration process using custom liquid classes.

The pipette tip is positioned 27 mm above the bottom of the well, corresponding to the area just above the interface with the melted paraffin layer. Prior to aspiration, 500 µL of warm air is slowly aspirated and dispensed to condition the tip and minimize paraffin solidification within the tip. A defined volume of melted paraffin (500–600 µL, depending on the cycle) is aspirated with tracking enabled. The liquid handler initially aspirates warm air, but due to tracking, will subsequently aspirate the melted paraffin. Aspiration is performed slowly (10 µL/s) to mitigate issues associated with viscous liquids and to prevent premature solidification of paraffin within the tip. For example, aspirating 600 µL at 10 µL/s requires 60 seconds. Please note that some buffer will also be aspirated as the tip does not stop aspirating past the paraffin layer. Tips containing paraffin are immediately discarded after each aspiration step.

### 2.4. Labware and Consumables

The selection of labware was guided by several criteria: compatibility with robotic gripper arms, suitability for centrifugation and cooling, accessibility for liquid handler arms, efficient heat transfer for reliable heating, prevention of spillover during shaking, and the ability to track samples (e.g., via barcodes). For this study, Azenta 1.0 mL 2D Barcoded Storage Tube (68-0702, Azenta Life Sciences) were selected due to their widespread use in long-term cryogenic storage and their 2D barcodes, which enable sample tracking. A significant challenge was identifying a suitable adapter that met requirements for robotic handling, centrifugation, and efficient heat transfer. Existing tube adapters with flat bottoms were suboptimal for heat transfer with the BioShake, and the loose fit of tubes limited shaking speeds. To address this, Axygen 2 mL deepwell plates (P-DW-20-C, Corning Life Sciences) were repurposed as tube adapters. The large well diameter accommodates Azenta tubes, and existing BioShake adapters (Adapter-NUNC® & Axygen® deep well 96/2.0 ml, 2016-1151) for these plates provide efficient heat transfer, as the heating elements envelop the wells.

### 2.5. Protein quantity from the comparison of the two cohorts

The comparative proteomic analysis demonstrated a high degree of overlap between the standard and automated tissue processing methods. A total of 8,305 proteins (96.1%) were identified by both approaches, while 47 proteins (0.5%) were uniquely detected using the standard method and 288 proteins (3.3%) were exclusively identified using the automated workflow. Furthermore, the automated method yielded a higher overall number of identified proteins with lower variability, as indicated by the reduced standard deviation in protein counts compared with the standard processing method (6146±860 vs. 4941±1091, p-value < 0.0001) (**Figure 6, A**). The principal component analysis (PCA) demonstrated substantial overlap between the two groups, indicating a high degree of similarity in their overall proteomic profiles. No clear separation or distinct clustering pattern was observed along the first two principal components (PC1: 5.1%, PC2: 3.8%), suggesting comparable molecular characteristics between the analyzed samples (**Figure 6, B**). Moreover, the higher efficiency of the automated method was also confirmed by pairwise comparison analyses of each individual sample (**Figure 6, C**). The correlation heatmap demonstrated a strong overall similarity between samples processed using the standard and automated tissue processing methods. Hierarchical clustering analysis revealed highly correlated protein expression profiles across the majority of samples, indicated by the predominantly intense red coloration throughout the heatmap (**Figure 6, D**), that automated tissue processing preserves proteomic patterns comparable to those obtained with the standard workflow. These findings further support the reproducibility and reliability of the automated method for downstream proteomic analyses.

**Figure 6.**
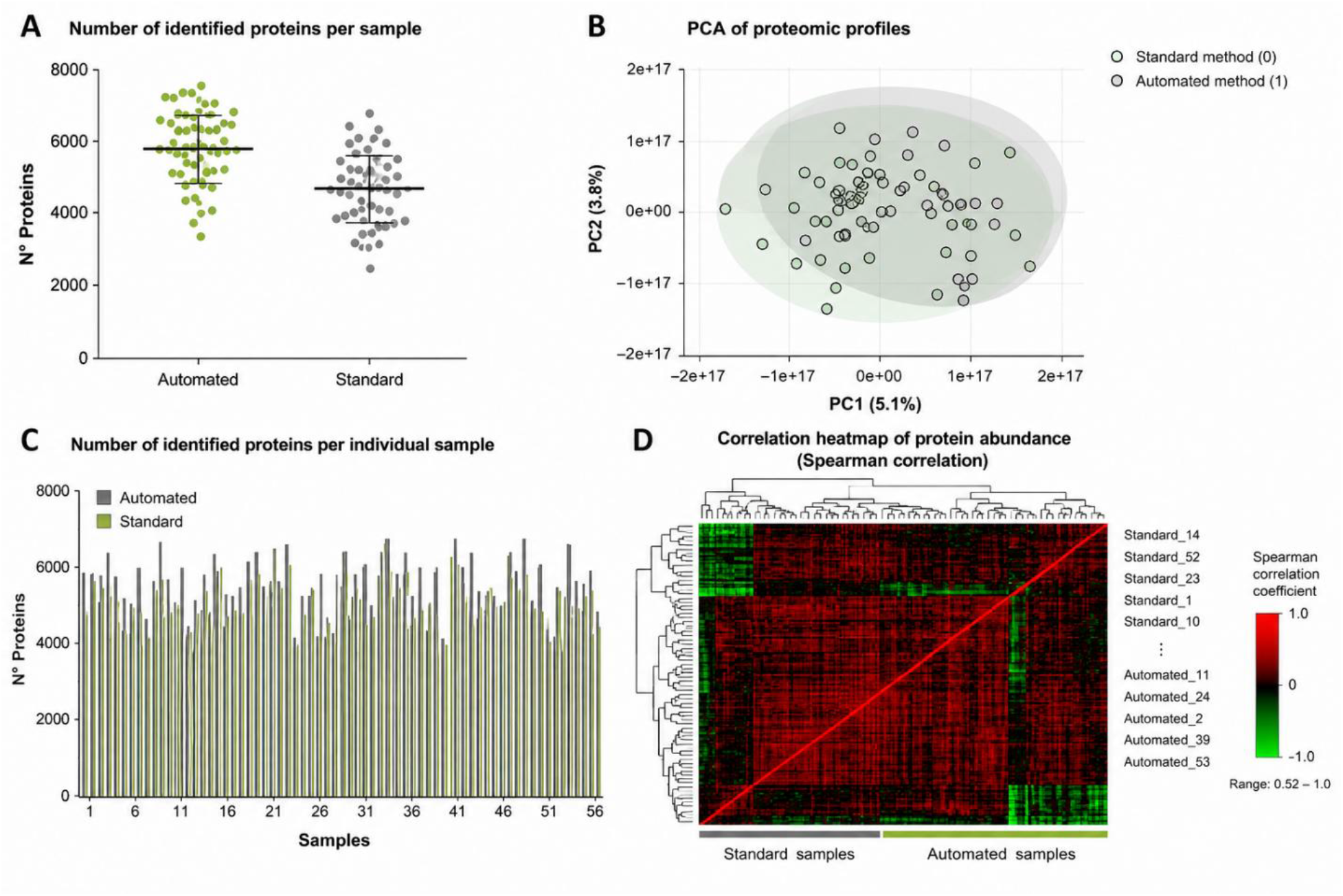
Comparative proteomic evaluation of standard and automated tissue processing methods. **(A)** represents a scatter plot showing the number of identified proteins in samples processed with automated and standard workflow. **(B)** highlights a PCA plot illustrating the clustering and distribution of samples processed by the two methods, indicating substantial overlap and overall similarity in proteomic profiles (0=standard method, 1=automated method). **(C)** Bar graph comparing the number of identified proteins across individual samples processed by the automated and standard methods. **(D)** Correlation heatmap with hierarchical clustering demonstrating strong overall similarity and high correlation between proteomic profiles generated by both tissue processing approaches. Color intensity reflects the Spearman rank correlation coefficient, ranging from 0.52 (green) to 1 (red). Pairwise correlation coefficients among all samples indicate high overall agreement between methods. Hierarchical clustering was performed based on Spearman correlation distance.

### 2.6. Molecular changes between the two subgroups

Proteomic analysis comparing primary melanoma samples with and without histological regression revealed significant differences in protein expression and pathway enrichment. A total of 97 proteins were upregulated and 226 proteins were downregulated in melanomas with regression. Pathway enrichment analysis demonstrated that proteins involved in mitochondrial and translational processes were significantly enriched in samples with regression, whereas proteins related to the innate immune system and complement pathways were predominantly downregulated. These results suggest that histological regression in primary melanoma is associated with distinct alterations in metabolic and immune-related biological processes (**Figure 7**).

**Figure 7.**
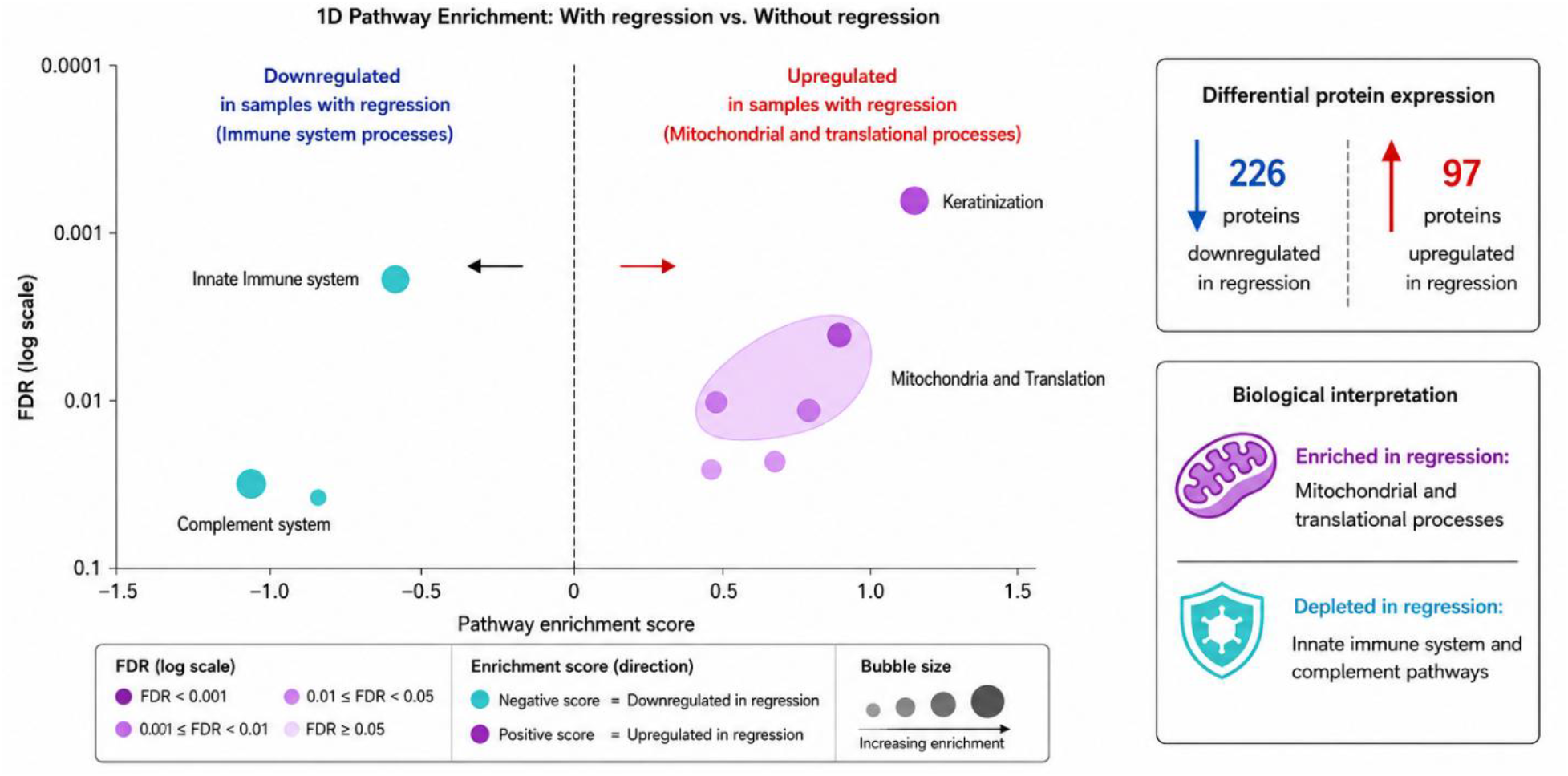
Differential proteomic and pathway enrichment analysis of primary melanoma samples with and without regression. The 1D enrichment comparing samples with and without regression. The presence of regression was associated with an upregulation of mitochondrial-related pathways and a downregulation of immune system pathways. The x-axis represents pathway enrichment score, while the y-axis indicates false discovery rate (FDR).

Importantly, beyond validating the workflow, the proteomic analyses demonstrated the robustness and reproducibility of the automated platform, confirming its ability to generate high-quality proteomic data with extensive protein coverage across melanoma tissue specimens. Furthermore, the analysis identified several candidate biomarkers and molecular features of potential biological relevance. These observations warrant further validation in independent cohorts and follow-up studies to confirm their reproducibility and evaluate their potential clinical significance.

## 3. DISCUSSION AND CONCLUSIONS

To better understand the mechanisms of melanoma progression or regression patient stratification is needed. For this purpose, the process of hundreds of thousands of samples from FFPE-derived repositories with consistent quality is essential. Automated tissue deparaffinization as a game changer enables high-throughput sample preparation rapidly leading to the acceleration of melanoma diagnostics, and supporting decision-making in urgent tumor cases.

FFPE tissues represent one of the most valuable resources for retrospective biomarker discovery and precision oncology, however, several technical limitations continue to affect proteomic performance. Among these, incomplete paraffin removal remains a major bottleneck, as residual paraffin may interfere with protein extraction, enzymatic digestion, and peptide ionization, ultimately reducing peptide yield and increasing analytical variability [9,10]. Furthermore, conventional solvent-based deparaffinization protocols may contribute to protein loss during repeated organic solvent exchange steps, particularly in laser microdissected or low-input melanoma samples where tissue availability is limited [11]. Formaldehyde-induced cross-linking additionally complicates efficient protein recovery by limiting peptide accessibility and extraction efficiency [9,10].

The critical next step is the industrial-scale implementation and full automation of the platform, enabling standardized, high-throughput workflows that deliver highly reproducible spatial tumor profiling and clinically actionable molecular readouts at true single-cell resolution. This capability represents a key technological frontier for the next generation of spatial biology platforms. We are advancing this development through our technology program in collaboration with Tecan ©, as well as the implementation of the tissue solubiliation technology be Covaris ©, establishing a fully automated, end-to-end processing workflow. The platform converts solid tumor tissue through a standardized, multistep process into liquid-phase samples optimized for downstream high-resolution protein expression analysis.

The translational relevance of such standardization is particularly important in dermatopathology and melanoma research, where large biobanked FFPE cohorts provide unique opportunities for molecular stratification and biomarker discovery. Our findings demonstrate that optimized and automated deparaffinization workflows can substantially improve the quality and reproducibility of proteomic analyses performed on FFPE melanoma tissues. In the present study, the automated workflow resulted in the identification of a higher number of proteins with lower technical variability compared with the conventional standard method, while maintaining highly comparable global proteomic profiles. These observations emphasize the importance of pre-analytical tissue preparation as a critical determinant of downstream molecular data quality in translational melanoma research.

Patient stratification based on the identification or prediction of tumor behavior in primary melanomas is fundamental to guiding appropriate therapeutic decision-making. In the present study, we compared predominantly thin, early-stage primary melanomas (AJCC 8th edition stage I–II) stratified by the presence or absence of histological regression. In line with our previous spatial proteomic investigations of early primary melanomas, mitochondrial activity emerged as a key molecular feature associated with more aggressive tumor behavior [14]. Aggressive melanoma phenotypes have been shown to exhibit recurrent activation of oxidative phosphorylation (OXPHOS), mitochondrial translation, the tricarboxylic acid (TCA) cycle, and RNA polymerase III-mediated transcriptional programs, with these features being particularly prominent in BRAF V600E-mutant tumors and cases associated with distant metastatic progression [14].

The relationship between histological regression and upregulated mitochondrial pathways remains incompletely understood. However, it is well established that regression reflects an active host inflammatory response, in which tumor-infiltrating lymphocytes and macrophages, both of which require mitochondrial metabolism playing a central role in supporting immune-cell function within the tumor microenvironment [15]. This mitochondrial dependence of immune effector cells may, at least in part, explain the paradoxical upregulation of mitochondrial pathways observed in regressing tumors. Furthermore, histological regression has been identified as a favorable prognostic factor in stage I and II cutaneous melanoma [16], lending clinical relevance to its molecular characterization.

In our cohort, histological regression in primary melanoma was associated with enhanced mitochondrial and translational activity alongside reduced activation of innate immune and complement pathways, reflecting distinct and potentially interrelated metabolic and immune alterations. These findings suggest that the proteomic landscape of regressing melanomas captures an active immune-metabolic crosstalk, which may underlie the more favorable clinical course observed in this subgroup.

In this context, our results support the implementation of heat-assisted, solvent-minimized, and automated tissue processing strategies to overcome these technical barriers. The strong overlap observed between the standard and automated workflows, together with the improved protein yield and lower variability achieved by automation, suggests that standardized deparaffinization may enhance both analytical robustness and reproducibility in large-scale melanoma proteomics. Importantly, the automated pipeline also demonstrated compatibility with downstream digital pathology workflows, which are increasingly integrated into modern precision oncology platforms.

## 4. METHODS

### 4.1. Patient samples

The study was carried out in strict accordance with the Declarations of Helsinki and was approved by the national-level Ethics Committee (Hungarian Scientific and Research Ethics Committee of the Medical Research Council, BM/15061-1/2023). Formalin-Fixed Paraffin-Embedded tissue (FFPE) tissue samples from the primary tumors of patients diagnosed with malignant melanoma were obtained from the Department of Dermatology and Allergology, University of Szeged. A total of 59 patients treated with primary cutaneous melanoma were included in the study. In two cases, there was no tumor content in the sample, and in three cases, the proteomic analysis was unsuccessful, therefore, these five samples were excluded from the statistical analysis. Based on the 54 patients, the mean age at primary melanoma diagnosis was 61.8 years (±18.1). The cohort consisted of 34 male and 20 female patients. The mean Breslow thickness was 1.33 mm (±2.4). Clinical staging at diagnosis was determined according to the 8th American Joint Committee on Cancer staging system. Most patients were diagnosed with predominantly early-stage melanoma, including stage IA (n=35) and stage IB (n=7). Less frequent stages included IIA-IV (n=12). Regarding histological subtype, superficial spreading melanoma (SSM) was the most common histotype (n=40), followed by SSM with vertical growth (n=2), nodular melanoma (NM) (n=4), lentigo maligna melanoma (LMM) (n=4), and other melanoma subtypes (n=4). Histological regression was present in 21 tumors and absent in 33 tumors. Ulceration of the primary tumor was observed in 19 cases, whereas 35 tumors were non-ulcerated. Overall survival (OS) was defined as the time from the diagnosis of the primary melanoma to death from any cause or the last documented follow-up. Disease-free survival (DFS) was defined as the time from primary melanoma diagnosis to the first documented melanoma recurrence or the last disease-free follow-up. Progression-free survival (PFS) was defined as the time from primary melanoma diagnosis to documented disease progression. Patients who remained event-free at the end of follow-up or were lost to follow-up before experiencing an event were treated as right-censored at their last documented follow-up. Survival analyses were performed using STATANow (version; BE/19.5, StataCorp LLC, College Station, TX, USA).

### 4.2. Sample processing and peptide/protein identification based on previous, manual techniques

Sample processing for proteomics involved a series of standardized steps to ensure high-quality protein extraction and digestion for whole tissue sections (WTS). For WTS, deparaffinization was performed using EnVision (1 mL of reagent with 10 min incubations at 95°C, followed by 15 min centrifugations at 15000 g at 18°C) until samples were paraffin-clear. A protein extraction buffer containing 5% SDS, 25 mM DTT, and 100 mM TEAB (pH 8.0) was added to the sliced tissues, incubating for one hour at 99 °C with shaking (500 RPM). Protein extraction was followed by sonication with a Bioruptor (40 cycles, 15 s on, 15 s off, at 4 °C) to enhance protein solubilization. Proteins were quantified either using the NanoDrop or the Pierce 660 nm Protein Assay (Thermo Scientific) supplemented with ionic detergent compatibility reagent (Thermo Scientific).

Protein digestion was conducted using S-Trap technologies (Protifi), with 96-well plates used for WTS samples. In brief, proteins were alkylated by adding IAA to a final concentration of 40 mM and incubated in the dark at room temperature for 30 minutes, followed by the addition of phosphoric acid (1.2%) and 6x final volume of S-Trap binding buffer (90% MeOH, 100 mM TEAB). Samples were loaded onto the filters with short centrifugations (1000 g), followed by 3 washing steps with the S-Trap binding buffer (each step using 200 uL). Proteins were digested with trypsin diluted in digestion buffer (50 mM TEAB) at a 1:25 enzyme-to-protein ratio at 37 °C. The enzyme addition was divided in three steps, followed by a 2-hour incubation in between each. Peptides were eluted in three steps, using 80 uL of the following: digestion buffer, 0.2% formic acid, and 50% ACN/ 0.2% formic acid. Each step was finalized with a short centrifugation for peptide elution (1000 g). Peptides were dried down in a centrifugal evaporator and resuspended in 2% ACN/0.1% TFA. The resulting peptide mixtures were quantified and stored at -80 °C until mass spectrometry (MS) analysis. Quantitative proteomics was performed, utilizing the Q-Exactive HF-X (Thermo) mass spectrometer and a data-independent acquisition (DIA) method for enhanced proteome coverage. This workflow ensured optimal protein extraction, digestion, and quantification, facilitating high-resolution proteomic analysis for molecular characterization of tumor heterogeneity.

### 4.3. Proteomic analysis (nLC-MS/MS)

The nano-LC MS/MS analysis was performed on an Ultimate 3000 HPLC coupled to a Q Exactive HF-X mass spectrometer (Thermo Scientific). Each sample (∼1 µg) was loaded onto a trap column (Acclaim1 PepMap 100 pre-column, 75 µm, 2 cm, C18, 3 mm, 100 Å, Thermo Scientific) and then separated on an analytical column (EASY-Spray column, 50 cm, 75 µm i.d., PepMap RSLC C18, 2 mm, 100Å, Thermo Scientific) using solvent A: 0.1% formic acid in water and solvent B: 0.1% formic acid in ACN, at a flow rate of 300 nL/min and a column temperature of 60°C. Peptide separation was achieved using a non-linear gradient over 100 minutes, beginning with an increase from 4% to 26% solvent B over 85 minutes, followed by a ramp to 50% B over the subsequent 15 minutes. Data were acquired using a recently implemented variable window data-independent acquisition (DIA) method [17].

### 4.4. Quality control of the MS analysis

Quality control measurements were introduced to assess the performance of LC-MS/MS systems. A protein digest from HeLa cells (Pierce HeLa Protein Digest Standard, Thermo Fisher Scientific) mixed with a standard peptide mixture (Pierce Peptide Retention Time Calibration Mixture) was used as a QC sample and measured every tenth LC-MS/MS analysis. This allowed monitoring of the peak width, retention time, base peak intensity, number of MS/MS, Peptide-Spectrum matches (PSMs), and number of peptides and proteins identified.

### 4.5. Data analysis

The generated proteomic raw data was processed in DIAnn and analysed by encompassing several stages to ensure comprehensive and accurate interpretation of the results. Initially, the raw data generated from whole tissue sections (WTS) were processed using DIAnn, an automated software suite designed for data-independent acquisition (DIA) proteomics. DIAnn employs deep neural networks and advanced quantification strategies to enhance the identification and quantification of proteins. In this analysis, the ‘match-between-runs’ (MBR) feature was utilized to align and match features across multiple runs, thereby increasing the depth of protein identification.

### 4.6. Data and Code Availability

The mass spectrometry-based proteomic data from the prospective, postmortem cohorts and the cell lines were deposit in PRIDE consortium. They were also listed in the Key Resources table. Raw proteomic datasets were processed independently using DIA-NN in directDIA mode, with identical parameter settings across both cohorts for quatitative analysis within the study. Spectral identification was carried out by searching against the UniProt human reference proteome. We applied a label-free quantification strategy, enforcing a 1% false discovery rate (FDR) for both peptide and protein identifications. The resulting data matrices were imported into R (version 4.4.2) for subsequent preprocessing. Protein abundance values were log2-transformed and then subjected to median centering at the protein level, and aligning each cohort’s distributions around the global median abundance. Technical replicates (multiple MS runs per sample) were consolidated by averaging protein abundance values. To mitigate batch effects between cohorts, we applied the remove BatchEffect function from the limma package (v3.62.2).

### 4.7. Digital pathology

Formalin-fixed, paraffin-embedded (FFPE) sections stained with hematoxylin and eosin (H&E) were digitized to enable high-resolution histopathological analysis. Expert pathologists annotated specific regions of interest including the aforementioned different structures of malignant melanoma regions, tumor infiltrating lymphocyte regions using whole-slide imaging. Tumor annotations were performed using QuPath v0.6.0 [18] and the SlideViewer (3DHISTECH Ltd., Budapest, Hungary) digital pathology platform were used.

## Author Contributions Statement

Jessica Guedes: Writing – review & editing, Visualization, Methodology, Investigation, Formal analysis. Leticia Szadai: Writing – review & editing, Methodology, Formal analysis, Data curation. Nicole Woldmar: Writing – review & editing, Investigation, Formal analysis, Data curation. Elisabet Wieslander: Writing – review & editing, Supervision, Conceptualization. Krzysztof Pawłowski: Writing – review & editing, Supervision. A. Marcell Szasz: Writing – review & editing, Supervision, Formal analysis, Data curation. Henriett Oskolas: Resources, Formal analysis, Data curation. Roger Appelqvist: Writing – review & editing, Supervision, Resources, Conceptualization. Johan Malm: Writing – review & editing, Supervision, Resources, Project administration, Funding acquisition, Conceptualization. Istvan Balazs Nemeth: Writing – review & editing, Supervision, Project administration, Investigation, Conceptualization. Jeovanis Gil: Writing – review & editing, Writing – original draft, Visualization, Supervision, Funding acquisition, Formal analysis, Data curation, Conceptualization. Gyorgy Marko-Varga: Writing – review & editing, Writing – original draft, Supervision, Resources, Project administration, Funding acquisition, Conceptualization.

## Declaration of Generative AI Use

During manuscript preparation, the authors used ChatGPT (OpenAI, GPT-5.1 [19]) to assist with language editing and improve clarity. All AI-supported text was carefully reviewed and revised by the authors, who assume full responsibility for the final content of the manuscript.

## Data Availability Statement

Proteomic data generated in this study have been deposited in the ProteomeXchange Consortium via the PRIDE partner repository. The accession number will be provided upon submission.

